# Tumor Spheroids Layered in an Imageable Cancer Environment (T-SLICE): a novel *in vitro* platform to study tumor biology

**DOI:** 10.1101/2022.10.08.511443

**Authors:** Morgan Pugh-Toole, Nicholas Dawe, Andrew Smith, Jeanette E. Boudreau, Brendan M. Leung

**Affiliations:** Department of Pathology, Dalhousie University; Beatrice Hunter Cancer Research Institute, Dalhousie University; School of Biomedical Engineering, Dalhousie University; Department of Microbiology & Immunology, Dalhousie University; Department of Applied Oral Sciences, Dalhousie University

## Abstract

Cancer treatment is shifting towards precise medicine informed by tumor genetics and structural features. In recent years, it has become increasingly recognized that patient tumors—even those of the same tissue origin—can differ substantially between patients and respond differently to treatment. Given this, it is necessary to design therapies that target the heterogeneity of tumors. When investigating novel therapeutics in the laboratory, conventional cell culture models do not adequately recapitulate this heterogeneity and therefore may not accurately represent drug responses. Recent advances in miniaturized organ-on-a-chip models have been able to generate more complex microenvironments for *in vitro* studies. However, many of these models do not resemble the scale of clinically relevant tumors and pose a high barrier to use because they are technically complex. To facilitate mechanistic studies of the tumor microenvironment (TME), we designed T-SLICE, a chip made using commercially available elastomers and designed to fit in a standard 6-well plate. This simple 3-D tumor model incorporates microfluidic principles into a fully customizable TME, wherein cells drive the formation of biochemical gradients akin to those observed within a real tumor. In T-SLICE, spheroids are seeded atop a monolayer of fibroblasts situated between two closely spaced coverslips (300-700 µm). The restrictive gap height limits the permeation of oxygen (O_2_) and hinders the removal of carbon dioxide (CO_2_) and metabolic waste, which leads to the generation of tumor-like hypoxic gradients. We demonstrate that T-SLICE establishes cell-driven oxygen gradients leading to the formation of a hypoxic core, with further impacts on cellular viability, mitochondrial membrane potential (MMP), and proliferation. T-SLICE cultures can be imaged live or fixed and stained for immunohistochemistry (IHC). These features of T-SLICE make it an accessible and faithful model of a tumor’s heterogeneity and open the possibility for more faithful testing of novel therapeutics in the context of a realistic TME.

## Introduction

Tumors are dynamic networks of tumor and stromal cells that together form a microenvironment with gradients of nutrients and oxygen^1^. Within the tumor microenvironment (TME) across various cancer histologies, the relative compositions of stromal and tumor cells vary, with impacts on tumor biochemistry, prognosis, and response to treatment^2^. The TME is highly dynamic and diverse, and biochemical factors within the TME allow cancer cells to persist and metastasize into healthy tissues. There are many different modes of studying the TME in 2-D or 3-D formats, however many existing models fall short of faithfully recapitulating these complexities^1, 3^. The emergence of microfluidic tumor models has allowed for the recapitulation of some tumor biochemistry; however, these are often complex and out of reach for cancer biology laboratories, and thus not implemented for cancer research^4–6^. In this work, we set out to create a simple, accessible, yet representative 3-D culture system that could be used to study the TME, and ultimately better inform its biology and the development of cancer therapies.

In tumors, hypoxia (low oxygen) is common at the tumor’s center as cancer growth outpaces vascularization^7–10^ . Acute and chronic hypoxic responses lead to the accumulation of reactive oxygen species (ROS) that can drive DNA damage and apoptosis in healthy cells^10^. Over time, the accumulation of irreparable DNA mutations gives rise to apoptosis resistance and advancing malignancies^11^. The rapid proliferation of cancer cells makes them metabolically demanding; this generates biochemical gradients that vary with tumor size, blood supply, and composition^12, 13^. Without sufficient mechanisms for nutrient delivery and waste clearance, compounded by hypoxia, cells at the tumor’s core evolve differently from those at its periphery. Tumor cells adapt, with manifestations of decreased proliferation and oxidative respiration, increased accumulation of mutations and ROS, and resistance to apoptosis^14, 15^. These layers of biochemical and tumoral diversity drive the selection of subsets of cancer cells that may confer resistance to treatment^14^.

Promising preclinical pharmaceutical and immunotherapy candidates demonstrate a low success rate in phase III clinical trials, with a cumulative failure rate of approximately 97%^16, 17^. Likely, these studies reflect that most tumors are only partially sensitive to therapeutic interventions because their heterogeneity is underrepresented in existing preclinical models. Over the past two decades, the applications of microfluidic technologies within microscale models have allowed for *in vitro* simulation of crucial biophysical features of solid tumors including chemical gradients, matrix stiffness, hydrodynamic fluid flow, and mechanical stresses ^18^. Various iterations of channel^19^ and multilayer^20^ microfluidic devices allow for the precise arrangement of tumor, stromal and immune cells within a defined microenvironment, while simultaneously imposing biochemical gradients. Moreover, the biochemical gradients driven by cell metabolism within diffusion-limiting microfluidic designs allow the development of a hypoxic core as established by cells within native tumors^21^. This was recently demonstrated in a channel-based microfluidic device with a thin gap culture chamber^22^. This model demonstrated the establishment of a hypoxic core driven by cell metabolism, with consequences on chemotherapy sensitivity as a function of oxygen tension^22^.

This demonstrates that these next-generation microfluidics may be better equipped to inform treatments of complex tumors.

The widespread adoption of microfluidic models for cancer therapy testing is hindered by the accessibility of these devices to biology laboratories. For example, the reliance on hydrophilic capillary forces to passively load cells into gradient-based microfluidics necessitates plasma or chemical activation and fundamentally limits the size of the device (and hence the tumor that can be modelled). In pre-assembled devices, pre-bonding of the device before cell loading eliminates the possibility of depositing cells, extracellular matrices and reagents in designated patterns with bioprinting techniques, so their spatial organization cannot be controlled. The arrangement, proximity and communication between cells each predicts tumor progression^23, 24^, so conventional pre-bonded microfluidic devices with closed channels cannot fully recapitulate important histological aspects of tumors^25, 26^.

To enable high-content and mechanistic studies that recapitulate the TME, we created <u>Tumor S</u>pheroids <u>L</u>ayered in an Imageable <u>C</u>ancer <u>E</u>nvironment (T-SLICE). T-SLICE incorporates principles of microfluidic design, is compatible with standard biology laboratories, and has tunable biochemical and cellular parameters that recreate features of the TME while remaining fully imageable. In T-SLICE, small spheroids (<100 cells) are seeded atop a fibroblast monolayer which is situated between two glass coverslips with a narrow vertical gap height (300-700 µm). The small gap space minimizes convective transport caused by bulk fluid flow, creating a diffusional resistance that is inversely proportional to its height. Within T-SLICE, cells situated closer to the edge of the parallel glass coverslips preferentially receive oxygen and nutrients. The arrangement of spheroids and fibroblasts within T-SLICE creates the topography, cellularity, and dimensionality characteristic of real tumors. The spheroids are small enough that intra-spheroid gradients are negligible, and the imageability limit of conventional spheroid culture is overcome by uncoupling tissue architecture from metabolically generated biochemical gradients. By enabling control of biochemical gradients independently from tumor tissue density, T-SLICE permits live cell imaging *in situ*, empowering high-content experiments to test therapeutic interventions. We demonstrate that the steepness of the hypoxic gradient is inversely proportional to the gap between the two coverslips and that with increasing exposure to hypoxia (i.e. proximity to the center of the T-SLICE chip), cellular proliferation, viability and mitochondrial membrane potential (MMP) decrease. Hence, in this simple model, into which cells, stroma and tumor cells can be arrayed as desired by the investigator to represent specific aspects of tumor histology, the hypoxic gradient and metabolic challenges with which it is associated are created. We thereby expect that T-SLICE will enable research that is better predictive of interactions and therapeutic responses in real tumors, accelerating the development of new treatments toward clinical trials.

## Materials & Methods

### 3-D mold design and printing

T-SLICE design and construction are depicted in **Figs 1A, and 1B**. T-SLICE chips were designed using Autodesk® Fusion 360^TM^ Computer-Aided Design (CAD) software as ’.stl’ files, then converted to ’.ctb’ files using Chitubox® 3-D printing preprocessing software. Designs were 3-D printed using an ELEGOO Mars 2 Pro MSLA 3-D Printer (ELEGOO, Shenzhen. China). Molds were created from Standard LCD UV-Curing Photopolymer Rapid Resin (ELEGOO, Shenzhen. China) in 0.05mm layers, each cured by 6 seconds of UV light exposure (405nm). Resin molds were washed in isopropanol (20 min) and UV-cured (405 nm, 30 min) (ELEGOO Mercury Plus Washing and Curing Machine, Shenzhen. China). Uncured resin monomers were extracted from molds by washing thrice in 95% ethanol for 24 h with agitation.

**Fig 1:**
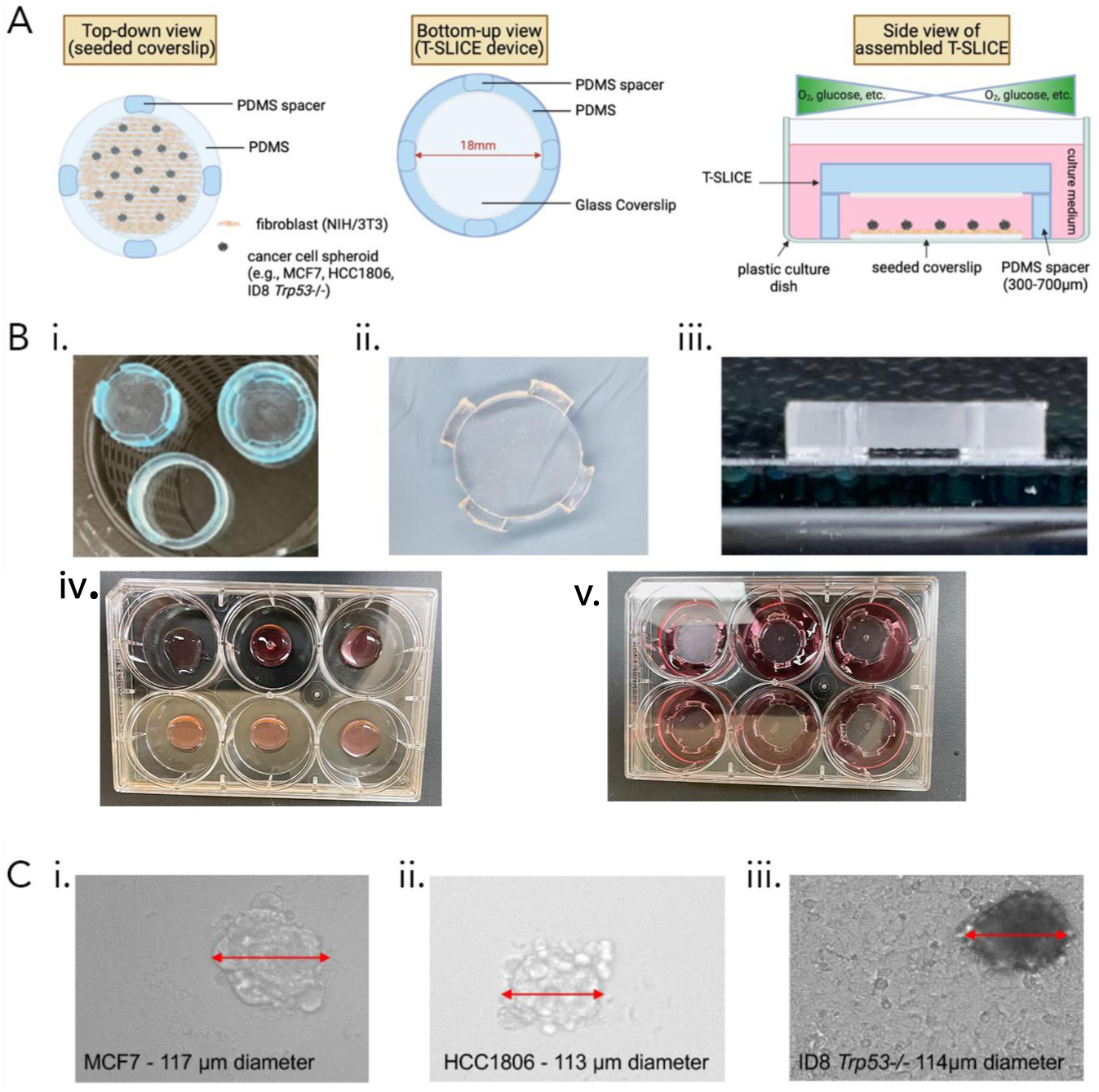
T-SLICE Schematic, Design & Assembly. **(A)** Top, bottom and side-view schematic of T-SLICE demonstrating the placement of chips and resulting biochemical gradients. (B) photograph of (i) two-part T-SLICE mold (shown both assembled, top right and in the component ring and insert pieces (bottom and top left); (ii) top-down and (iii) side view of assembled T-SLICE; (iv) Top-down view of T-SLICE cultures assembled in a six-well plate without and (v) with a T-SLICE chip added on top. (C) photographs depicting tumor cell line spheroids atop fibroblast monolayers imaged through a T-SLICE chip: (i) MCF7; (ii) HCC1806; (iii) ID8 *Trp53*^-^

T-SLICE chips were prepared by pouring Polydimethylsiloxane (PDMS) (10:1 elastomer to curing agent, Sylgard 184™ Silicone Elastomer Kit, Corning) into the resin molds, degassed in a vacuum chamber for 15 min, then cured (60°C for 24h) and removed using forceps. An 18mm circular glass coverslip was covalently bonded to the underside of the chip by plasma oxidation (1-2 min). Before use in cell culture, T-SLICE chips were sterilized in 70% ethanol and exposed to UV light for 30 min.

### Cell Culture

The NIH/3T3 (fibroblast), MCF-7 and HCC1806 (breast cancer) cell lines were obtained from the American Type Culture Collection (ATCC), and the ID8 *Trp53^-/-^* (murine ovarian cancer) cell line was generously gifted by Dr. Madhuri Koti (Queen’s University). Cells were maintained in a humidified incubator at 37°C and 5% CO_2_. MCF-7 and NIH/3T3 cells were maintained Dulbecco’s Modified Eagle Media (DMEM; Gibco) supplemented with 10% fetal bovine serum (FBS; Gibco) and 1% antibiotic-antimycotic (AA;100 IU/mL penicillin, 0.1 mg/mL streptomycin and 0.25 μg/mL amphotericin-B, Gibco,). HCC-1806 cells were maintained in RPMI-1640 media (Gibco) supplemented with 10% FBS and 1% AA. ID8-*Trp53^-/-^* cells^12^ were maintained in DMEM supplemented with 10% FBS, 1% penicillin-streptomycin (PS; Wisent), and 1% Insulin Transferrin Selenite (ITS; Wisent Bioproducts).

### Spheroid culture and T-SLICE assembly

Tumor spheroids were formed using Perfecta3-D^TM^ 384-well hanging drop culture plates (3-D Biomatrix Inc. Ann Arbor, MI). Hanging drop plates were pretreated in 0.1% w/v Pluronic® F108 solution (Sigma-Aldrich) in distilled water, then rinsed with dH_2_O and UV-sterilized for 30 min prior to use. MCF-7 or HCC1806 and ID8-*Trp53-/-* were seeded in 25 µl droplets containing 50-100 cells, supplemented with 2.4 mg/mL MethoCel® A4M (Sigma-Aldrich) to create spheroids with diameters of <150 μM **(Fig 1C)**. To prevent spheroids from merging, droplets were seeded in a lattice pattern, yielding 192 spheroids for each 384-well hanging drop plate. Spheroids were maintained in standard cell culture conditions (37°C, 5% CO_2_). Following spheroid formation (24h for MCF-7; 48h for HCC1806 and ID8-*Trp53-/-*), spheroids were collected, centrifuged at 200 x *g*, and resuspended in 500 μL cell culture media, which was then used to replace the culture media atop the fibroblasts.

T-SLICE cultures were assembled in standard six-well cell culture plates. NIH/3T3 fibroblasts were cultured on autoclaved No. 1 18mm round coverslips (VWR) pretreated with fibronectin (10 μg/mL in PBS; Sigma-Aldrich) **(Fig 1B)**. The pretreated coverslips were seeded with NIH/3T3 fibroblasts overnight at 200,000 cells per coverslip (400,000 cells/mL) to reach 100% confluency, and then 192 spheroids were added dropwise to each coverslip. After overnight co-culture of the fibroblasts and spheroids, the T-SLICE chip was placed on top of the coverslips, and 2 mL of cell culture media was added at the sides of the T-SLICE chip to fill it by capillary action. Media was partially exchanged by replacing 500mL every 24 h. Control wells of co-cultures without T-SLICE chips were cultured in parallel to ascertain the impact of the chip on cellular processes. Zones of T-SLICE were defined as middle, intermediate and peripheral zones, representing concentric rings of 0-3 mm, 3-6 mm or 6-9 mm diameter, respectively.

### EF5 characterization of hypoxic responses

To characterize hypoxic cellular responses, we cultured T-SLICE in the presence of 100 mM of the EF5 Hypoxia Detection Kit according to the manufacturer’s instructions (Sigma-Aldrich). EF5 (2-(2-Nitro-1H-imidazol-1-yl)-N(2,2,3,3,3-pentafluoropropyl) acetamide), is a 2-nitroimidazole compound that is reduced in hypoxic cells to form adducts at the cell membrane that can be fluorescently labelled with a fluorophore-conjugated anti-EF5 antibody^13^. Cell culture media was supplemented with 100 μM EF5 and was replenished every 24 hours. Cultures were assessed at 6, 12, 24, or 48 h time points. After the T-SLICE chip was removed, cells were fixed in 4% paraformaldehyde (PFA) in PBS overnight at 4°C in the dark. Cells were then washed and prepared for imaging, using the anti-EF5 (clone ELK3-51-Cy3) at 5 mg/mL in 1% BSA/PBS (Sigma Aldrich). Imaging was performed on the EVOS^TM^ FL Auto 2 Cell Imaging System, and analysis was conducted using ImageJ. Imaging was completed using the EVOS^TM^ Light Cube, RFP 2.0 (Ex: 542 nm; Em: 593nm).

### Assessment of Cellular proliferation using EdU

Cell proliferation in T-SLICE zones was analyzed using an EdU (5-ethynyl-2’-deoxyuridine) Cell Proliferation Kit (Click-iT^TM^ EdU Cell Proliferation Kit for Imaging, Alexa Fluor^TM^ 488 dye: Invitrogen). Cell culture media was supplemented with 10 mM EdU which was replenished every 24 h. After 72 h, T-SLICE chips were removed, and the cells were fixed with 4% PFA in PBS for 15 min at RT and then stained according to the manufacturer’s instructions. Imaging was performed using the EVOS^TM^ Light Cube, GFP 2.0 (Ex: 482nm; Em: 524nm).

### Assessment of cell viability using Calcein-AM and Ethidium homodimer-1

Cell viability was assessed using Calcein-AM (ThermoFisher Scientific) to label live cells and ethidium homodimer-1 (EthD; Invitrogen) to label dead cells. At specified timepoints, T-SLICE chips were removed, and then cells were washed with PBS and stained with a PBS solution containing 1 μM Calcein-AM and 1 μM ethidium homodimer-1 for 30 min at rt in the dark. Cells were washed twice with PBS and imaged using the EVOS^TM^ Light Cube, GFP 2.0 (Ex: 482 nm, Em: 524 nm) and EVOS^TM^ Light Cube, RFP 2.0 (Ex: 542 nm, Em: 593 nm).

### Mitochondrial Membrane Potential measurement with MitoView 633

MitoView^TM^ 633 (Biotium) was used to measure mitochondrial membrane potential (MMP) according to the manufacturer’s recommendations. Cell culture media was supplemented with 100 nM MitoView 633 (Biotium) for the duration of T-SLICE culture. T-SLICE cultures were imaged live every 24 h using the EVOS^TM^ Light Cube, Cy5 2.0 (Ex: 635 nm, Em: 692 nm).

### Statistical analysis

Statistical analysis of data was performed using GraphPad Prism 9 (GraphPad Software). Two-way analysis of variance (ANOVA) with Tukey’s multiple comparisons test was performed to assess multi-group comparisons and α was set to 0.05. *P*-values are represented as follows:

\**p<0.05; **p<0.01; ***p<0.001; ****p<0.0001*.

## Results

### Conceptualization, design and development of T-SLICE

We sought to design a model that recapitulates the biochemical properties of the solid tumor amenable to use in biology laboratories. We considered chip geometry, cell growth surface area, and distance between the two parallel glass coverslips as variables that would influence biochemical gradients. Mitochondrial health, cell proliferation and vitality are all linked to tumor cell diversification and oxygen availability, so we prioritized the generation of hypoxia in our design. We hypothesized that cellular metabolism by a co-culture of spheroids and stromal cells would generate an oxygen gradient with a hypoxic core in a chip where diffusion of oxygen and nutrients through the media would be restricted to the lateral plane. Other key features considered in this iterative process included ease of use and compatibility with live cell imaging and endpoint IHC.

Throughout the design process, we tested several prototypes, including different shapes and sizes and elected to use a circular chip for its symmetry. We elected to assemble a chip that was comprised of a standard 18 mm round coverslip bonded on the underside of a polydimethylsiloxane (PDMS) chip of 24 mm diameter. Below the chip, a second 18 mm round coverslip was situated onto which cells would be cultured.

We first conducted a series of simulations using COMSOL Multiphysics® v. 6.2. (www.comsol.com, COMSOL AB, Stockholm, Sweden) to predict the range of heights that would promote hypoxia and thus, the useful range of T-SLICE chip heights. We modelled the generation of hypoxia as a function of time (**Fig 2A**), and at three gap heights (300, 500, 700 mm, **Fig 2B**), predicting the radial O_2_ concentration from the periphery to the center of T-SLICE (radius = 9 mm). We assumed the outer edge (peripheral) O_2_ concentration to be 6.7 mg/L (0.21 mol/m^3^), which is the dissolved oxygen saturation in water at sea-level^10^. To predict how O_2_ tension would change from the periphery to the center of T-SLICE at 37°C and by metabolism of cells on the bottom coverslip, we assumed that no O_2_ diffusion was occurring through the top and bottom glass layers, and set the bottom layer of the model as a surface reaction layer to simulate a constant oxygen consumption rate per cell of 6 x 10^-17^ mol/cell/s, scaled to the average density of a confluent cell monolayer (2.67x10^4^ cells/cm^2^). This resulted in an overall consumption rate of 1.6 x 10^-8^ mol/(m^2^ s) ^27^.

**Fig 2:**
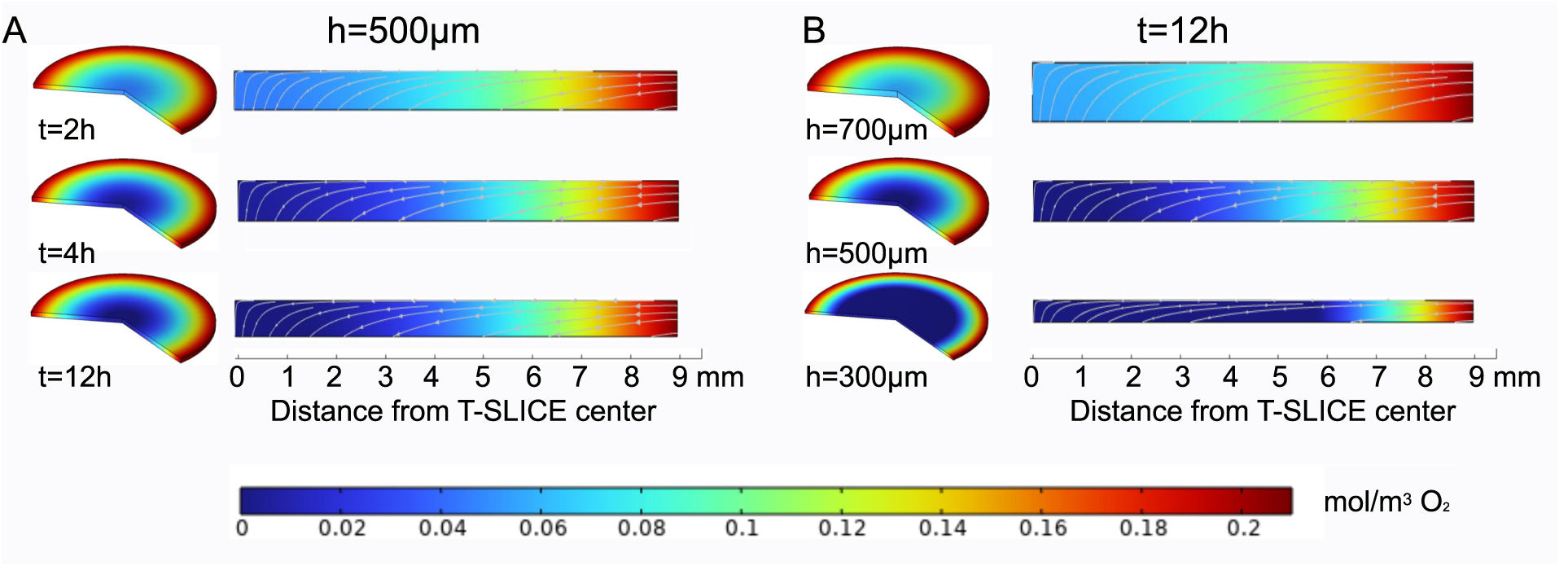
Computational Simulations of Oxygen Gradients in T-SLICE. Expected radial oxygen concentration gradient within the T-SLICE as a function of (A) time (500 mm gap height) and (B) gap heights (at 12 hours). Oxygen diffusion through the culture media under T-SLICE was simulated using COMSOL Multiphysics, where a time-dependent transport of diluted species study was constructed. A rectangle was created in the 2D axisymmetric space dimension with the left side of the rectangle as the axis of symmetry. The length was set to the radius of the T-SLICE (9 mm), and the height was varied to match the different gap heights (300, 500, 700 mm). The material in the space between the top and bottom was modelled as water at 37 °C to simulate culture media: the diffusion coefficient was set to 3 x 10-9 m^2^/s, and the initial concentration of oxygen was set to the dissolved oxygen saturation point of 6.7 mg/L (0.21 mol/m^3^) based on literature values. Data are presented as a heatmap from low (0 mol/m^2^, dark blue) to high (0.2 mol/m^2^, dark red) oxygen concentrations.

In our simulations, both time and gap height impacted the onset and degree of hypoxia. Over a simulation period of 24h, the model showed that a steady state oxygen gradient was reached after 12 h in the 500 µm T-SLICE chip, where an anoxic core of 3 mm diameter was predicted in the center (**Fig 2A**). The study was then repeated for T-SLICE with 300 µm and 700 µm gap heights. Steady-state results (T= 12 h) predicted an increase in anoxic core diameter to 11.2 mm when the gap height was reduced to 300 µm. Inversely, when the gap height is increased to 700 µm the anoxic core is eliminated with a center minimum oxygen concentration equal to 0.0549 mol/m^3^ (**Fig 2B**).

### Functional testing of the T-SLICE model

To test the validity of our simulations and move toward a working T-SLICE model *in vitro*, we assembled co-cultures of fibroblasts and spheroids under T-SLICE chips (**Figs 1A and 1B**). Fibroblasts are one prominent cell type found in the TME of many tumors, therefore we chose to incorporate them in T-SLICE to represent the stromal compartment of tumors. We used NIH/3T3 embryonic murine fibroblasts as the gradient-driving stromal monolayer in all T-SLICE co-cultures described. The NIH/3T3 cells were grown to 100% confluency on 18 mm coverslips. Once confluent, tumor spheroids derived from hanging drop culture were seeded on top of the fibroblast monolayer. We tested a range of cell seeding densities and found that 2x10^5^ NIH/3T3 cells per coverslip established a confluent monolayer in 24 h, onto which tumor spheroids could be supported without direct contact with the coverslip and associated deformation due to spheroid adhesion spreading **(Figs 1B and 1C)**. In parallel, we optimized cancer cell seeding densities in hanging drop culture to create spheroids with a diameter <150 µm. We chose this limit to minimize the generation of intra-spheroid biochemical gradients and to ensure that the oxygen availability to the spheroid is dependent only on its position under the T-SLICE chip. The small size of the spheroid further allows microscopic imageability and quantification of biochemical changes in real-time (**Fig 1C**).

### T-SLICE culture generates sustained hypoxic gradients

Hypoxia is a hallmark feature of the TME of solid tumors. It results from limited oxygen availability, especially in regions distal to the blood supply^28^. Our simulations predicted that hypoxia would occur most prominently in the middle of the T-SLICE and exhibit relative normoxia at its periphery. To validate this functionally, we identified three concentric zones for analysis: middle (M, 0– 3 mm from center), intermediate (I, 3– 6 mm from center), and periphery (P, 6– 9 mm from center). Using EF5 staining, we measured hypoxia in T-SLICE after 6, 12, 24 and 48 h of culture. At all three gap heights tested, increasing time resulted in greater hypoxia among ID8 *Trp53-/-* cells (p<0.0001, **Fig 3**); this result was confirmed in two additional second cell lines, MCF7, and HCC1806 (**Supplementary Figs 1 and 2**) and consistent with our simulations (**Fig 1)**. For all cell lines and gap heights tested, EF5 fluorescence intensity was highest in the middle zone of T-SLICE and progressively decreased in the intermediate and peripheral zones. EF5 staining was observed as early as 6h and increased during the 48 h during which we studied cells in T-SLICE for this experiment (p<0.005 for all gap heights). Hypoxic gradients were most pronounced with the smallest gap height (300 µm) compared to the larger gap heights, consistent with our prediction of decreasing oxygen availability with restricted diffusion as gap height decreases (p<0.0001 at all timepoints).

**Fig 3:**
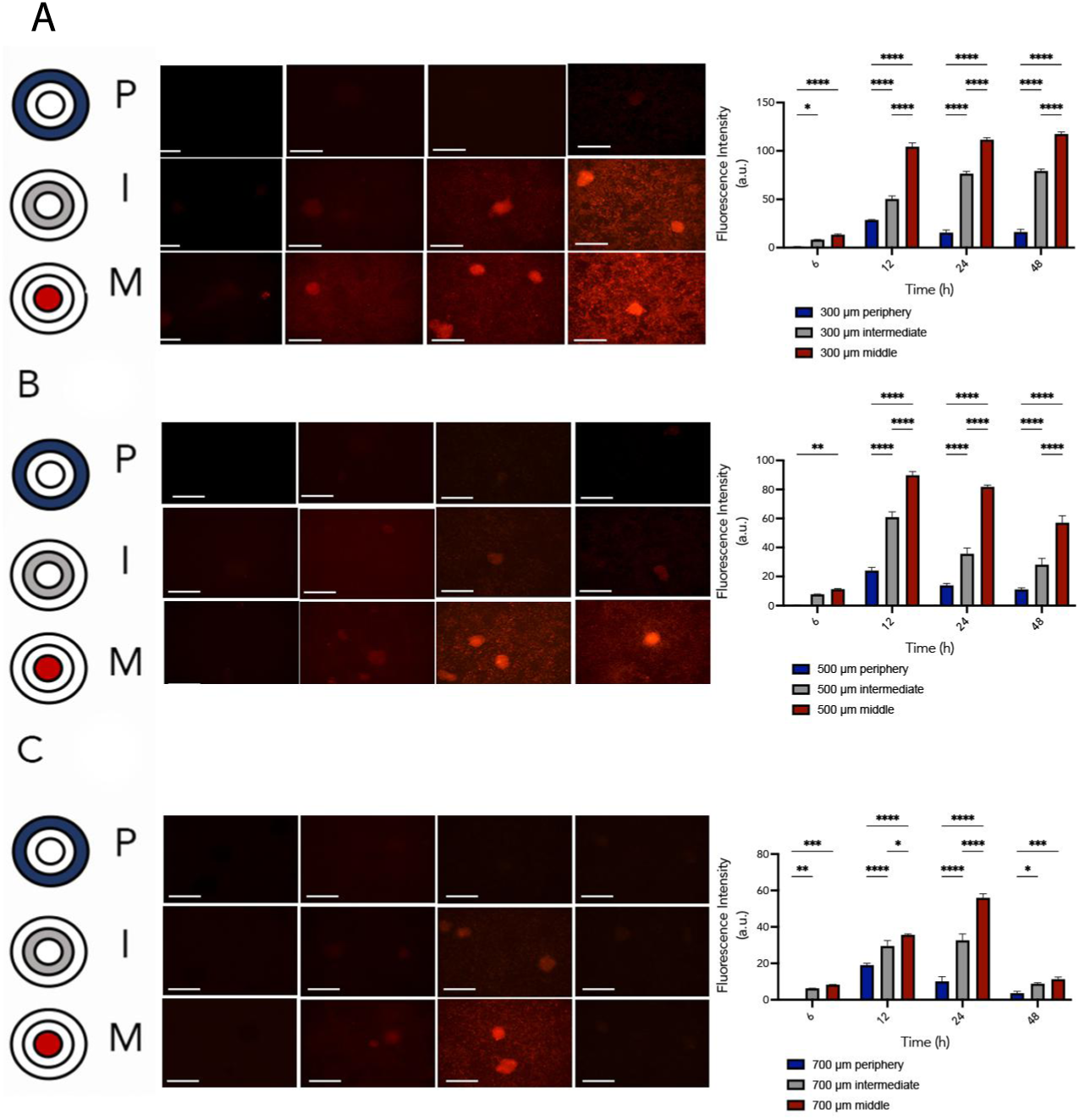
T-SLICE generates hypoxic gradients from the periphery to the center. *ID8 Trp53-/-* spheroids were seeded atop a fibroblast monolayer for 6, 12, 24 or 28h in a T-SLICE chip with EF5-containing media. Thereafter, cells were fixed and stained with anti-EF5 to ascertain hypoxic responses (A-C): photographs and fluorescence quantifications; (D-E) co-cultures under T-SLICE without the EF5 compound. P: periphery, I: intermediate, M: middle. A.u: arbitrary units. Scale bars represent 275mm. Data are representative of three technical replicates with measurements of fluorescence intensity in three fields of view for each zone (p<0.0001, error bars represent standard deviation (SD).

### Cell proliferation is diminished in hypoxic regions of T-SLICE

A direct impact of hypoxia in cancer cells is the reduction of proliferation to maintain viability^29^. We used the EdU assay to assess whether cellular proliferation is diminished in the hypoxic regions of T-SLICE. After 72h of culture, proliferation of tumor cells in spheroids and fibroblasts was most often observed in the peripheral zone of T-SLICE (p<0.0001, **Fig 4 and Supplementary Fig 3**). Newly divided MCF-7 or HCC1806 cells were not detected in the middle zone, at any gap height. In MCF7 cells, cell proliferation increased with higher gap heights (p<0.001), confirming that cellular proliferation and hypoxia are inversely related in T-SLICE.

**Fig 4:**
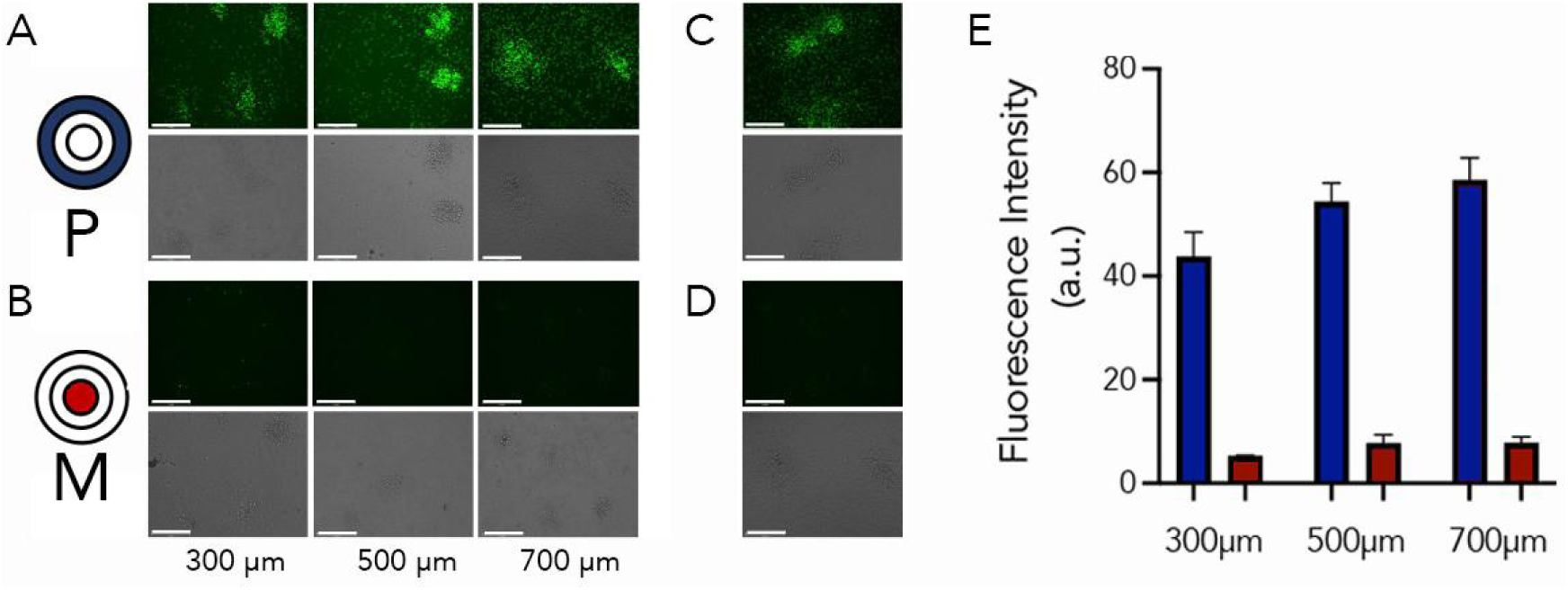
Cellular proliferation is halted in the middle region of T-SLICE. MCF-7 cells were seeded on a fibroblast monolayer in T-SLICE in the presence of EDU. After 72h incubation, cellular proliferation (EDU incorporation) was ascertained by fluorescence microscopy. (A) peripheral zone; (B) Middle zone; C) No T-SLICE control; D) unstained control; (E) quantification of fluorescence. a.u., Arbitrary units. Blue bars represent quantification from the peripheral zone; red bars represent quantification from the middle zone. Scale bars represent 275mm. Data are representative of three technical replicates with measurements of fluorescence intensity in three fields of view for each zone (p<0.0001, error bars represent SD).

### Cell viability is reduced in the hypoxic regions of T-SLICE

Concurrent with reduced proliferation, loss of cell viability is commonly observed in hypoxic regions of tumors due to deprivation of essential nutrients and oxygen^28^. To examine cellular viability within T-SLICE, we cultured cells for 72h and used Calcein-AM and ethidium homodimer-1 to stain in tandem to identify live (green) and dead (red) cells, respectively **(Fig 5)**. Cell viability remained nearly 100% in the peripheral zone of T-SLICE at all gap heights tested. In ID8 *Trp53^-/-^* spheroids cultured in 300 µm gap height T-SLICE, cell viability decreased to ∼70% and ∼20% in the intermediate and middle zones, respectively, but this diminution was nearly completely abrogated when gap heights were raised to 500 µm and 700 µm. Similar cellular viability trends were observed when we instead used MCF-7 cells **(Supplemental Fig 4).**

**Fig 5:**
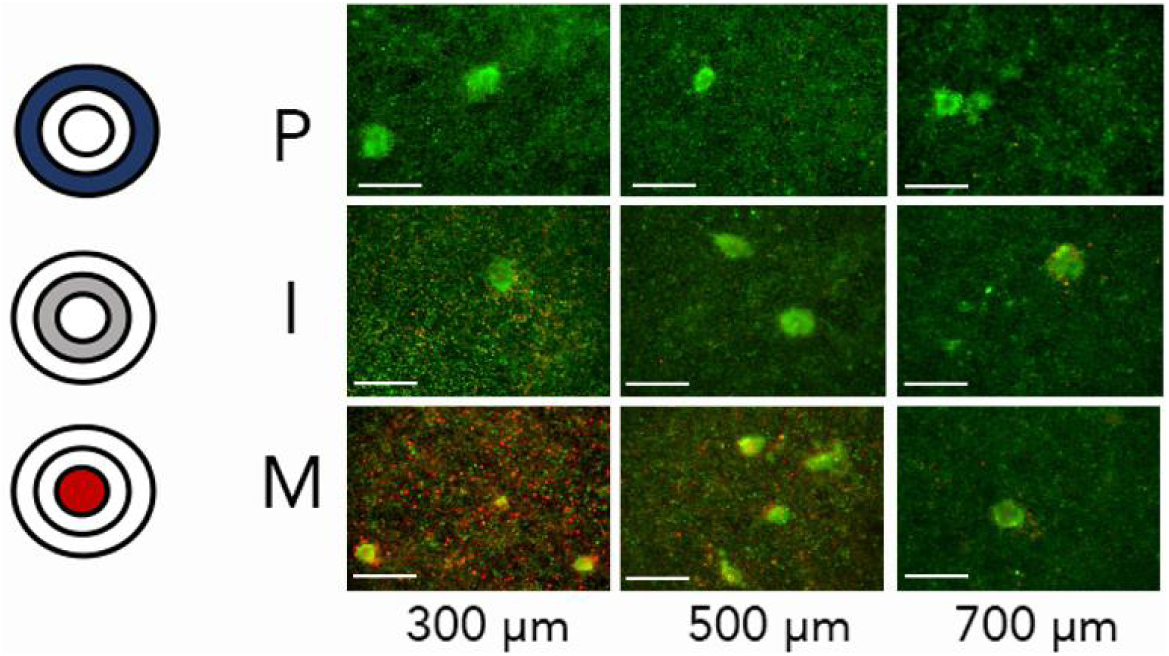
Cellular viability is lost in the most hypoxic regions of T-SLICE. ID8-Trp53^-/-^ spheroids were cultured in T-SLICE on a fibroblast monolayer for 72h, then stained with Calcein-AM and ethidium homodimer to visualize cell viability. Images are representative of three technical replicates and three fields of view for each T-SLICE zone. P, periphery; I, intermediate; M, middle.

### Hypoxia slowed proliferation and cell death in T-SLICE correspond to loss of mitochondrial membrane potential

Loss of mitochondrial membrane potential (MMP) is a characteristic early feature of apoptosis and can be brought about by hypoxic stress^29^. MitoView 633 was used to analyze the MMP of cancer cells cultured in T-SLICE for up to 48 h **(Fig 6 and Supplementary Fig 5)**. In both MCF-7 and ID8 *Trp53-/-* spheroids, MMP was significantly diminished in the intermediate zone, and further in the middle zone of T-SLICE compared with the periphery (p<0.001). This impact was significantly greater after 48 h (p<0.001). Altogether, our results indicate that T-SLICE creates biochemical gradients of hypoxia that correspond to loss of MMP, thereby impacting cellular viability and proliferation, recapitulating the features of native tumors.

**Fig 6:**
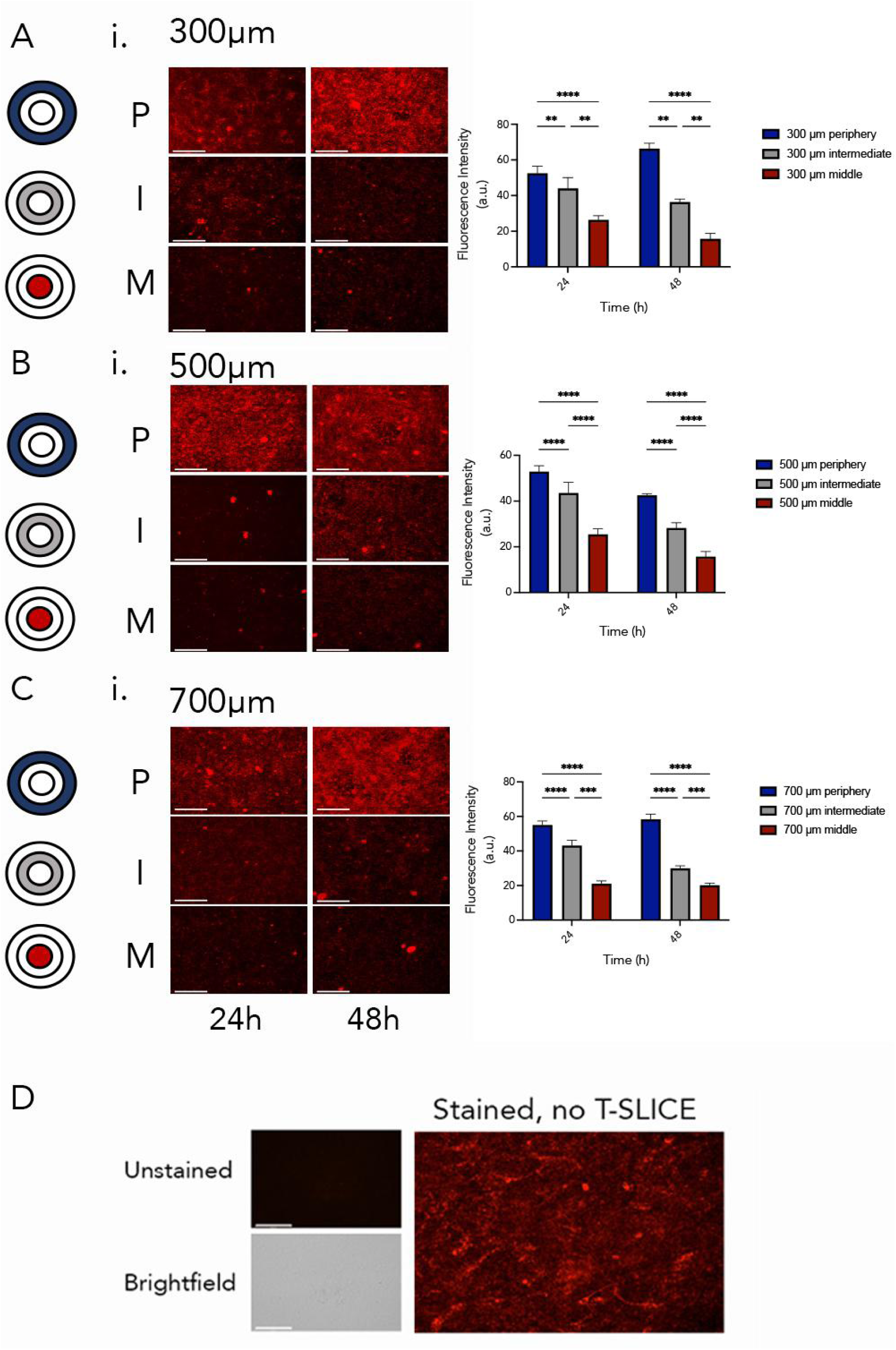
Mitochondrial membrane potential is reduced in T-SLICE. MMP was measured using MitoView 633 staining after 24 or 48h in MCF7 cells. Significant diminishment was observed in the intermediate and middle zones compared to the periphery (p<0.001) with greater impact after 48h (p<0.001). (A-C) each of the 300, 500 and 700 μM gap heights were tested. (i) representative images; and (ii) quantification are shown. (D) controls included unstained cells in T-SLICE, and stained cells with no T-SLICE. P, periphery (blue); M, middle (grey); I, intermediate (red); a.u., Arbitrary units. Scale bars represent 275mm. Data are representative of three technical replicates with measurements of fluorescence intensity in three fields of view for each zone (p<0.001 after 24h, p<0.0001 after 48h, error bars represent SD).

## Discussion

Hypoxia is a persistent feature of tumors that establishes much of their heterogeneity and fuels their potential progression and escape from therapeutic interventions^10, 15, 28, 30^. To catalyze cancer research, we created T-SLICE, a simple and effective *in vitro* system that uses a fibroblast monolayer seeded between impermeable top and bottom barriers to establish a cell-driven hypoxic microenvironment. Because the transparent chip is assembled atop a coverslip laden with cells, T-SLICE is fully compatible with live cell imaging. In its simplicity and flexibility, T-SLICE is highly amenable to use in biology laboratories and opens the possibility of testing new therapeutics in a model representative of a native tumor.

In our studies, we worked with three tumor cell lines that represented breast and ovarian cancer. With some inter-tumor variation, all tested cell lines generated the same biochemical trends: greater proximity to the middle of T-SLICE corresponded to a reduction in viability, proliferation, and MMP. Compared to cancer models that rely on extrinsically imposed oxygen gradients^31, 32^, a cell metabolism-driven gradient is more likely to recapitulate the TME, with flexibility for the investigator to tailor the model to a given tumor histology or characteristics. This is especially valuable given the wide range of oxygen tensions at normoxia within different tissue types^33^. Although we did not directly measure oxygen tension in T-SLICE, EF5 staining, cell viability and proliferation confirmed the generation of hypoxic gradients and their downstream consequences, and their behaviour is consistent with other cell-driven hypoxic cancer models^34, 35^. We confirmed that the area of the hypoxic zone in T-SLICE can be increased by reducing the chip gap height. A minimum gap height must be maintained to accommodate spheroids (<150 mm), but by increasing the diameter of T-SLICE, one could increase diffusion distances from the periphery to the middle and thus further increase the hypoxic area. By manipulating the gap height and T-SLICE diameters, we can mimic different TMEs and hypoxia levels, making T-SLICE representative of different-sized tumors or at different regions of a tumor with varying distances from blood vessels. By comparison, in spheroid culture, a gold standard practice in cancer research, an increase in hypoxic effect can only be achieved with more cells and larger spheroid diameters which will directly impair the ability to observe cell behaviors within^36^. Using T-SLICE, biochemical gradients can be uncoupled from tissue architecture, so it becomes possible to observe cellular events within a comparable TME to a much larger spheroid.

The use of animals as a preclinical screening model has been standard practice in drug development for decades, but the legislation of the FAD Modernization Act 2.0 in 2022^37^ encourages alternative model development, including organ-on-a-chip models such as T-SLICE, to improve the predictive power of preclinical studies and lower the cost of drug development. With increasing research into personalized and precision medicine approaches for cancer therapy, there is an urgent need to develop simple, reliable, high-content systems that can be widely adopted by biology research laboratories. Conventional 2-D culture and model organisms have been informative for understanding mechanisms behind disease processes; however, 2-D systems lack a tumor’s structural and biochemical characteristics. Model organisms, including xenografted or humanized mice, are excellent tools to study cancer in a complex microenvironment, but lack a human immune system, require extensive workup and monitoring^38^, and have much lower throughput compared to *in vitro* platforms, and can be quite heterogeneous.

Microfluidic platforms combined with 3-D culture systems allow a small number of explanted tumor, immune, and stromal cells to be grown and studied, but their complex setups make them out of reach for most biology laboratories^39^. Endpoint sample recovery can be challenging such as in sealed microfluidic chips, further impeding their use as a standard model to study cancer biology. In contrast, T-SLICE can be easily assembled using standard cell culture vessels (i.e., 6-well plates), which allows relatively high-throughput tumor studies with continuous monitoring capability in a broadly affordable and feasible manner. The transparency of T-SLICE combined with thin cell layers enables brightfield and fluorescent live-cell imaging. By simply lifting the T-SLICE chip with forceps, selected portions of cells or the entire culture can be harvested for endpoint histologic analysis using standard staining protocol, all without disrupting the overall spatial arrangement of cells.

An advantage that T-SLICE shares with microfluidic culture platforms is the low cell number needed to establish culture and biochemical gradients due to the small culture volume. We show that simple adjustments, such as changing the gap height of the chip, have predictable impacts on the hypoxic gradient; this is relevant because oxygen availability varies even within the same tumor^40, 41^. Presently, the hypoxia in T-SLICE is driven primarily by the fibroblast monolayer; however, relative proportions of the tumor, stromal and other cells should be carefully considered and included to reflect specific applications of T-SLICE.

In clinical practice, T-SLICE could be used to inform precision therapy. With fewer than 100,000 cells, patient “avatars” could be established in a recreated TME. Models using conventional spheroid culture would require several magnitudes more cells and would not be imageable. In T-SLICE, features and interventions can be studied with iteration in adjacent wells of the same plate to test, iterate, and combine treatments to identify the best one for each patient, and on a timeline consistent with informing care^40, 41^.

### Conclusions

T-SLICE is a novel *in vitro* tumor model that captures the biochemical features of the TME. With the shift away from mouse models, and toward the use of miniaturized organ-on-a-chip models for drug development pipelines, there is an unmet need for simple, accessible models. T-SLICE captures key aspects of tumor biochemistry while remaining amenable to techniques and equipment found in standard biology laboratories. By combining features of existing *in vitro* technologies, such as spheroid culture, and subjecting them to an environment that intrinsically imposes TME features, T-SLICE can meet the demand for accessible tumor model systems.

### Author contributions

The T-SLICE model and study was conceptualized by MPT, ND, AS, JEB, and BML; experiments were designed, conducted, and validated by MPT, ND, and AS. Data analysis was conducted by MPT, ND, AS, JEB, and BML. The manuscript was written by MPT, ND, AS, JEB, and BML, reviewed, edited, and approved by all authors.

## Acknowledgements

The authors thank Ms. Sarah Nersesian and Dr. Naeimeh Jafari for technical assistance and discussions on T-SLICE development. MPT and ND are trainees in the Cancer Research Training Program of the Beatrice Hunter Cancer Research Institute, with funds for MPT previously provided by the Bruce and Dorothy Rosetti Endowment through the Dalhousie Medical Research Foundation and funds for ND previously provided by the Helyer/Williams Studentship. MPT is funded by a CIHR Canadian Graduate Scholarship - Doctoral and a Scotia Scholars Scholarship. ND was supported by a Canadian Graduate Scholarship (Masters; NSERC) and a Genomics in Medicine Scholarship through the Dalhousie Medical Research Foundation. JEB is the Dalhousie Medical Research Foundation Cameron Cancer Scientist. We acknowledge the support of the Government of Canada’s New Frontiers in Research Fund (NFRF), [NFRFE2019-00415] for funding this research. Funding was additionally provided by Health Canada to Ovarian Cancer Canada in support of the OvCAN research initiative. We thank Dr. Madhuri Koti (Queen’s University) for the ID8-Trp53^-/-^ cell line.

## Data availability

All data generated or analysed during this study are included in this published article and supplementary data.

## References

1. B. Emon, J. Bauer, Y. Jain, B. Jung and T. Saif, Comput Struct Biotechnol J, 2018, 16, 279–287.

2. F. R. Balkwill, M. Capasso and T. Hagemann, Journal of Cell Science, 2012, 125, 5591–5596.

3. G. Imparato, F. Urciuolo and P. A. Netti, International Materials Reviews, 2015, 60, 297–311.

4. Y. Liu and H. Lu, Curr Opin Biotechnol, 2016, 39, 215–220.

5. J. Wu, H. Fang, J. Zhang and S. Yan, *J*ournal of Nanobiotechnology, 2023, 21, 85.

6. G. Velve-Casquillas, M. Le Berre, M. Piel and P. T. Tran, Nano Today, 2010, 5, 28–47.

7. Y. Li, L. Zhao and X. F. Li, Technol Cancer Res Treat, 2021, 20, 15330338211036304.

8. G. Chen, K. Wu, H. Li, D. Xia and T. He, Front Oncol, 2022, 12, 961637.

9. Z. Chen, F. Han, Y. Du, H. Shi and W. Zhou, Signal Transduction and Targeted Therapy, 2023, 8, 70.

10. V. Petrova, M. Annicchiarico-Petruzzelli, G. Melino and I. Amelio, Oncogenesis, 2018, 7, 10.

11. X. Jing, F. Yang, C. Shao, K. Wei, M. Xie, H. Shen and Y. Shu, Mol Cancer, 2019, 18, 157.

12. S. Romero-Garcia, J. S. Lopez-Gonzalez, J. L. Báez-Viveros, D. Aguilar-Cazares and H. Prado-Garcia, Cancer Biol Ther, 2011, 12, 939–948.

13. C. Navarro, Á. Ortega, R. Santeliz, B. Garrido, M. Chacín, N. Galban, I. Vera, J. B. De Sanctis and V. Bermúdez, Pharmaceutics, 2022, 14.

14. C. Carmona-Fontaine, M. Deforet, L. Akkari, C. B. Thompson, J. A. Joyce and J. B. Xavier, Proceedings of the National Academy of Sciences, 2017, 114, 2934–2939.

15. R. Abou Khouzam, K. Brodaczewska, A. Filipiak, N. A. Zeinelabdin, S. Buart, C. Szczylik, C. Kieda and S. Chouaib, Front Immunol, 2020, 11, 613114.

16. C. H. Wong, K. W. Siah and A. W. Lo, Biostatistics, 2018, 20, 366–366.

17. F. Spagnolo, A. Boutros, F. Cecchi, E. Croce, E. T. Tanda and P. Queirolo, BMC Cancer, 2021, 21, 425.

18. T. K. Ngan Ngo, C.-H. Kuo and T.-Y. Tu, Biomicrofluidics, 2023, 17.

19. H. Mollica, Y. J. Teo, A. S. M. Tan, D. Z. M. Tan, P. Decuzzi, A. Pavesi and G. Adriani, Biomaterials Science, 2021, 9, 7420–7431.

20. M. Virumbrales-Muñoz, J. Chen, J. Ayuso, M. Lee, E. J. Abel and D. J. Beebe, Lab on a Chip, 2020, 20, 4420–4432.

21. D. M. Cochran, D. Fukumura, M. Ancukiewicz, P. Carmeliet and R. K. Jain, Annals of Biomedical Engineering, 2006, 34, 1247–1258.

22. J. M. Oh, H. M. Begum, Y. L. Liu, Y. Ren and K. Shen, ACS Biomaterials Science & Engineering, 2022, 8, 3107–3121.

23. G. Corredor, X. Wang, Y. Zhou, C. Lu, P. Fu, K. Syrigos, D. L. Rimm, M. Yang, E. Romero, K. A. Schalper, V. Velcheti and A. Madabhushi, Clin Cancer Res, 2019, 25, 1526–1534.

24. D. E. Cohn, A. Forder, E. A. Marshall, E. A. Vucic, G. L. Stewart, K. Noureddine, W. W. Lockwood, C. E. MacAulay, M. Guillaud and W. L. Lam, Front Immunol, 2023, 14, 1275890.

25. S. Nersesian, R. J. Arseneau, J. P. Mejia, S. N. Lee, L. P. Westhaver, N. W. Griffiths, S. R. Grantham, L. Meunier, L. Communal, A. Mukherjee, A.-M. Mes-Masson, T. Arnason, B. H. Nelson and J. E. Boudreau, Frontiers in Immunology, 2024, 14.

26. F. Katou, H. Ohtani, Y. Watanabe, T. Nakayama, O. Yoshie and K. Hashimoto, Cancer Research, 2007, 67, 11195–11201.

27. C. H. Cho, J. Park, D. Nagrath, A. W. Tilles, F. Berthiaume, M. Toner and M. L. Yarmush, Biotechnol Bioeng, 2007, 97, 188–199.

28. V. Bhandari, C. Hoey, L. Y. Liu, E. Lalonde, J. Ray, J. Livingstone, R. Lesurf, Y. J. Shiah, T. Vujcic, X. Huang, S. M. G. Espiritu, L. E. Heisler, F. Yousif, V. Huang, T. N. Yamaguchi, C. Q. Yao, V. Y. Sabelnykova, M. Fraser, M. L. K. Chua, T. van der Kwast, S. K. Liu, P. C. Boutros and R. G. Bristow, Nat Genet, 2019, 51, 308–318.

29. X. Zhao, L. Liu, R. Li, X. Wei, W. Luan, P. Liu and J. Zhao, Med Sci Monit, 2018, 24, 8722–8733.

30. W. Zeng, P. Liu, W. Pan, S. R. Singh and Y. Wei, Cancer Lett, 2015, 356, 263–267.

31. L. Orcheston-Findlay, A. Hashemi, A. Garrill and V. Nock, Microelectronic Engineering, 2018, 195, 107–113.

32. H. Nam, K. Funamoto and J. S. Jeon, Biomicrofluidics, 2020, 14, 044107.

33. A. Carreau, B. El Hafny-Rahbi, A. Matejuk, C. Grillon and C. Kieda, J Cell Mol Med, 2011, 15, 1239–1253.

34. B. Mosadegh, M. R. Lockett, K. T. Minn, K. A. Simon, K. Gilbert, S. Hillier, D. Newsome, H. Li, A. B. Hall, D. M. Boucher, B. K. Eustace and G. M. Whitesides, Biomaterials, 2015, 52, 262–271.

35. D. Rodenhizer, T. Dean, B. Xu, D. Cojocari and A. P. McGuigan, Nature Protocols, 2018, 13, 1917–1957.

36. M. Roy, C. Alix, A. Bouakaz, S. Serrière and J. M. Escoffre, Pharmaceutics, 2023, 15.

37. P.-J. H. Zushin, S. Mukherjee and J. C. Wu, The Journal of Clinical Investigation, 2023, 133.

38. N. C. Walsh, L. L. Kenney, S. Jangalwe, K. E. Aryee, D. L. Greiner, M. A. Brehm and L. D. Shultz, Annu Rev Pathol, 2017, 12, 187–215.

39. J. H. Hammel, S. R. Cook, M. C. Belanger, J. M. Munson and R. R. Pompano, Annu Rev Biomed Eng, 2021, 23, 461–491.

40. G. J. Yoshida, J Hematol Oncol, 2020, 13, 4.

41. P. Voabil, M. de Bruijn, L. M. Roelofsen, S. H. Hendriks, S. Brokamp, M. van den Braber, A. Broeks, J. Sanders, P. Herzig, A. Zippelius, C. U. Blank, K. J. Hartemink, K. Monkhorst, J. B. A. G. Haanen, T. N. Schumacher and D. S. Thommen, Nature Medicine, 2021, 27, 1250–1261.

