## Supplemental file for "Tumor Spheroids Layered in an Imageable Cancer Environment (T-SLICE): a novel *in vitro* platform to study tumor biology"

Mailing Address: 5891 University Avenue, Dentistry Building, Room 5193, Halifax NS, B3H 1W2

**Supplemental Figures**

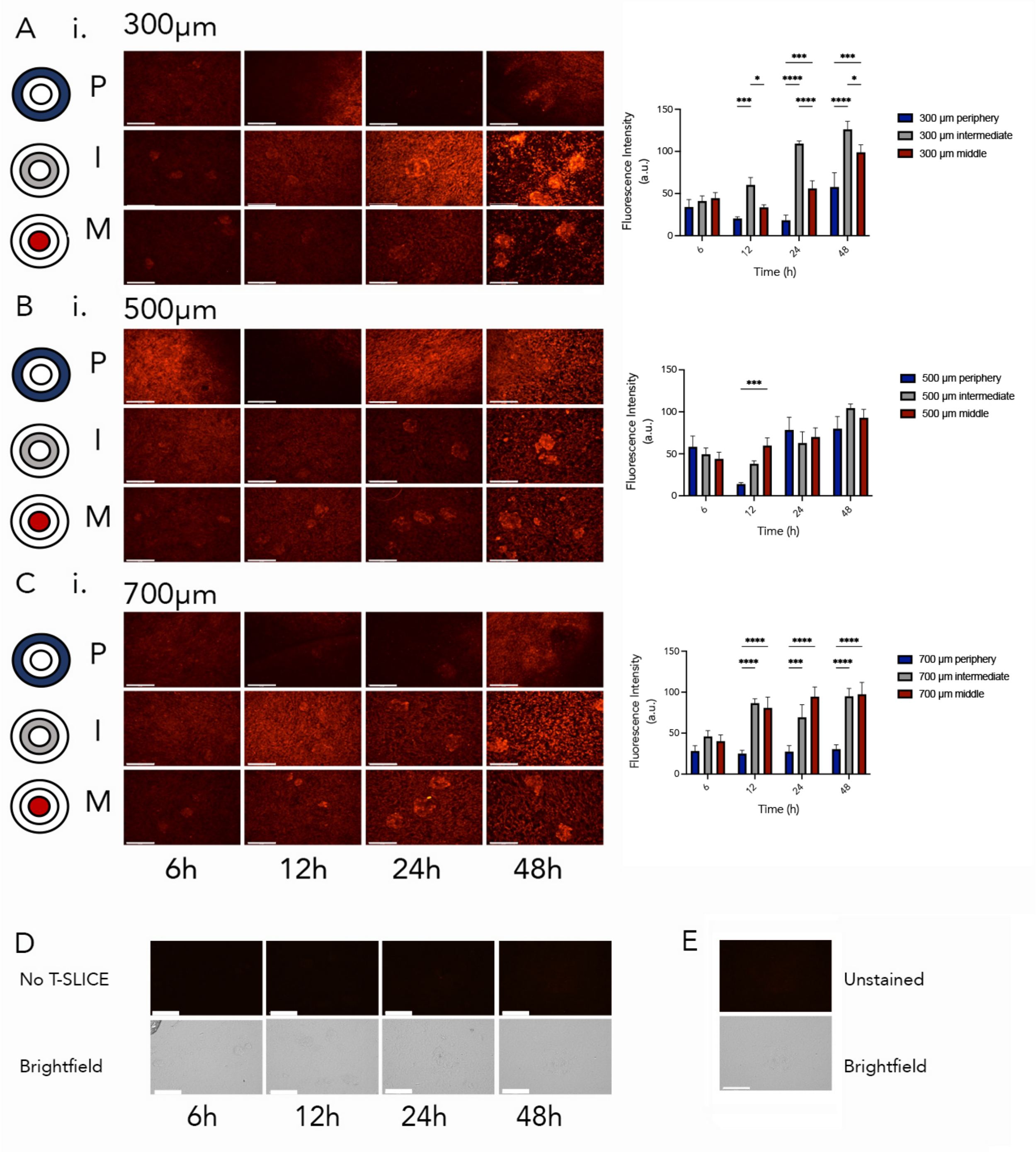


**S1 Fig. T-SLICE generates hypoxic gradients from the periphery to the center in MCF7 cells**. MCF7 spheroids were seeded atop a fibroblast monolayer for 6, 12, 24 or 48h. Thereafter, cells were fixed and stained with EF5 to ascertain hypoxic responses (A-C). (i) ) photographs of EF5

staining in designated T-SLICE culture regions and; (ii) corresponding fluorescence quantifications. (D and E) control wells either did not have a T-SLICE installed (D) or did not include dye (E). P, periphery (blue); I, intermediate (red); M, middle (grey); a.u., Arbitrary units. Scale bars represent 275 μm. Data are representative of three technical replicates with measurements of fluorescence intensity in three fields of view for each zone ($p<0.0001$, error bars represent standard deviation (SD)).

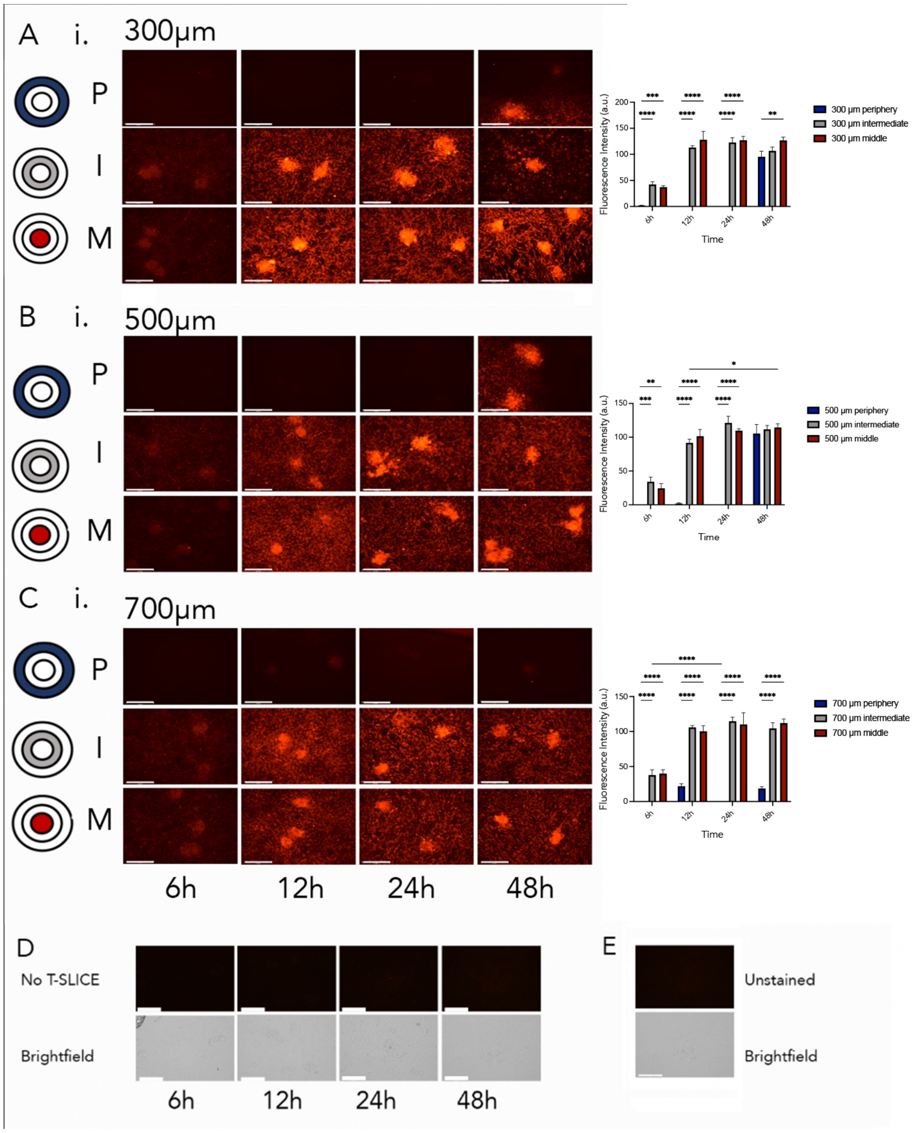


**S2 Fig. T-SLICE hypoxic gradients from the periphery to the center in HCC1806.** HCC1806 spheroids were seeded atop a fibroblast monolayer for 6, 12, 24 or 48h. Thereafter, cells were fixed and stained with EF5 to ascertain hypoxic responses (A-C). (i) photographs of EF5 staining in designated T-SLICE culture regions and; (ii) corresponding fluorescence quantifications. (D and E) control wells either did not contain dye (D) or did not have a T-SLICE chip installed I. P, periphery; I, intermediate; M, middle. Scale bars represent 275μm. Data are representative of three technical replicates with measurements of fluorescence intensity in three fields of view for each zone ($p<0.0001$, error bars represent SD).

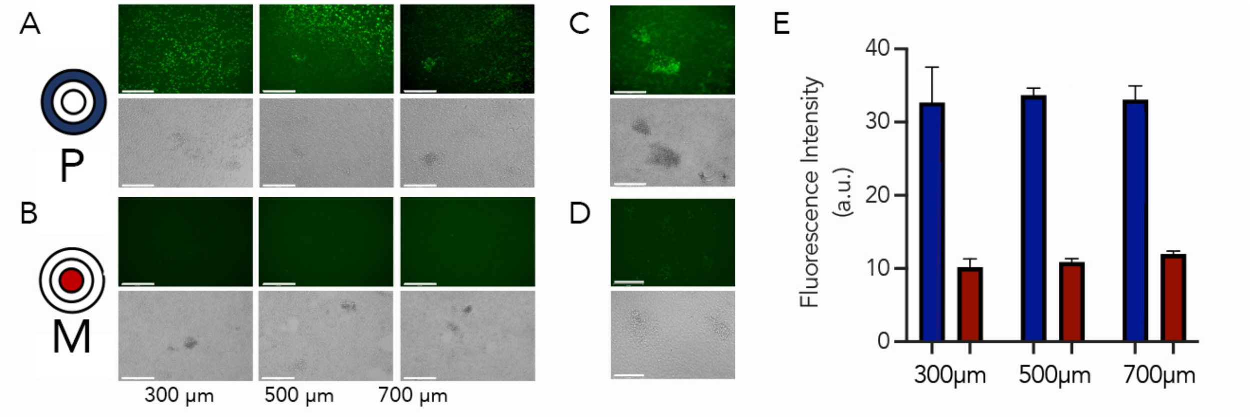


**S3 Fig. Cellular proliferation is slowed in HCC1806 cells in T-SLICE**. HCC1806 cells were seeded on a fibroblast monolayer in T-SLICE in the presence of EDU. After 72h incubation, cellular proliferation (EDU incorporation) was ascertained by fluorescence microscopy. (A) peripheral zone; (B) Middle zone; C) No T-SLICE control; D) unstained control; (E) quantification of fluorescence. a.u., Arbitrary units. Scale bars represent 275µm. Data are representative of three technical replicates with measurements of fluorescence intensity in three fields of view for each zone ($p<0.0001$, error bars represent SD).

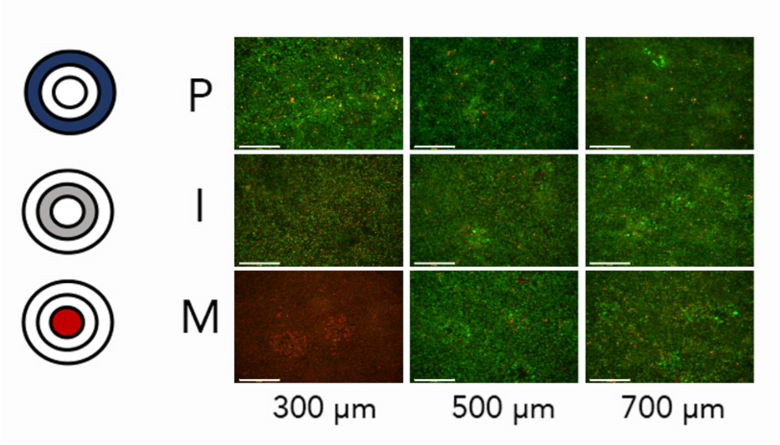


**S4 Fig: Cell viability is lost in the most hypoxic regions of T-SLICE.** MCF7 spheroids were cultured in T-SLICE on a fibroblast monolayer for 72h, then stained with Calcein-AM and ethidium homodimer to visualize cell viability. Images are representative of three technical replicates and three fields of view for each T-SLICE zone. P, periphery; I, intermediate; M, middle. Scale bars represent 275μm.

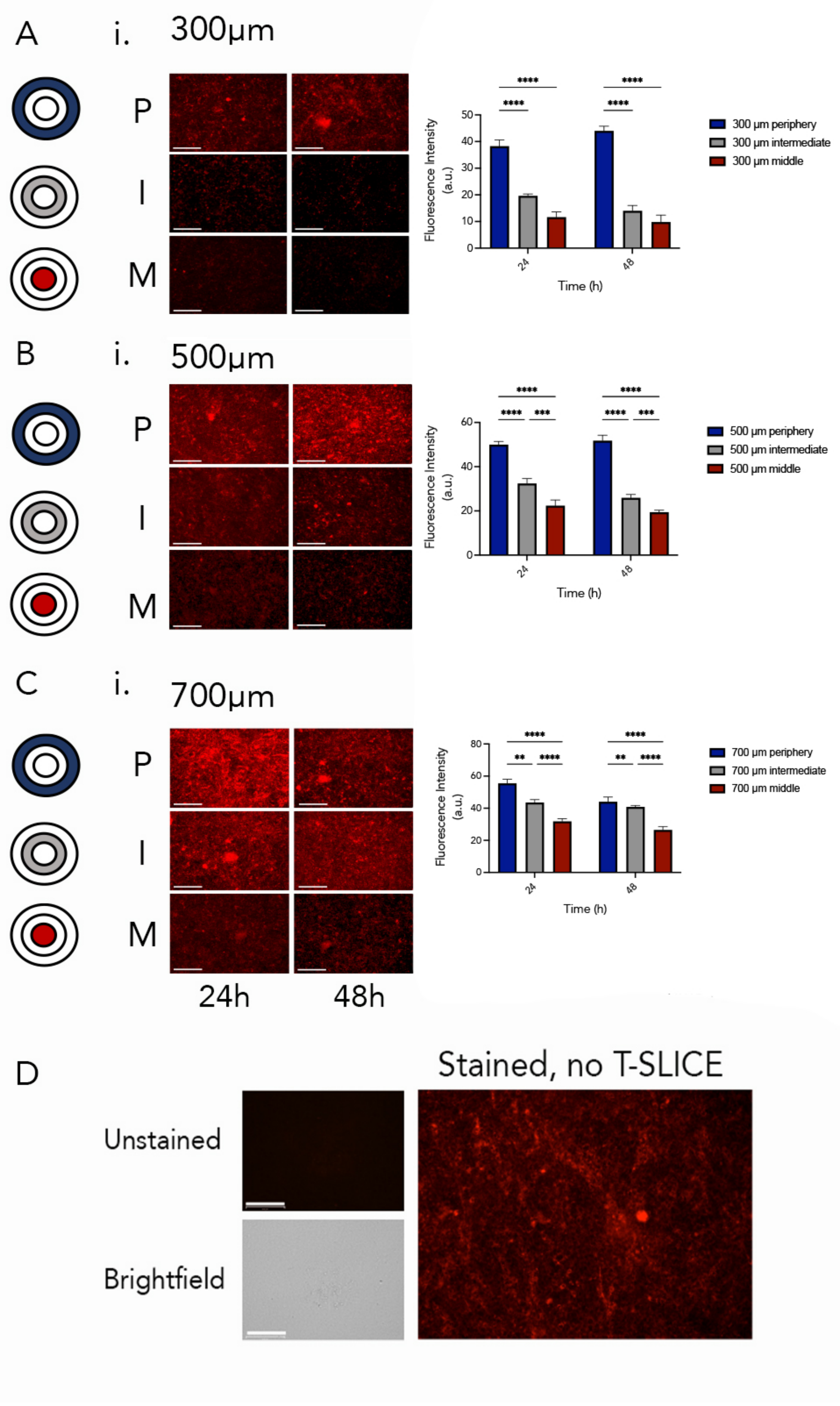
A
i.
300μm
P
I
M
Fluorescence Intensity (a.u.)
Time (h)
300 μm periphery
300 μm intermediate
300 μm middle
B
i.
500μm
P
I
M
Fluorescence Intensity (a.u.)
Time (h)
500 μm periphery
500 μm intermediate
500 μm middle
C
i.
700μm
P
I
M
Fluorescence Intensity (a.u.)
Time (h)
700 μm periphery
700 μm intermediate
700 μm middle
24h
48h
D
Stained, no T-SLICE
Unstained
Brightfield

**S5 Fig: MMP is reduced in T-SLICE in ID8-Trp53$^{-/-}$ cells.** MMP was measured using MitoView 633 staining after 24 or 48h in ID8-Trp53-/- cells. (A-C) each of the 300, 500 and 700 μm gap heights were tested. (i) representative images; and (ii) quantification are shown. (D) controls included unstained cells in T-SLICE, and stained cells with no T-SLICE. P, periphery (blue); M, middle (grey); I, intermediate (red); a.u., Arbitrary units. Scale bars represent 275 μm. Data are representative of three technical replicates with measurements of fluorescence intensity in three fields of view for each zone ($p<0.001$ after 24h, $p<0.0001$ after 48h, error bars represent SD).
